# Glioblastoma recurrence and the role of MGMT promoter methylation

**DOI:** 10.1101/317636

**Authors:** Katie Storey, Kevin Leder, Andrea Hawkins-Daarud, Kristin Swanson, Atique U. Ahmed, Russell C. Rockne, Jasmine Foo

## Abstract

Tumor recurrence in glioblastoma multiforme (GBM) is often attributed to acquired resistance to the standard chemotherapeutic agent temozolomide (TMZ). Promoter methylation of the DNA repair gene MGMT has been associated with sensitivity to TMZ, while increased expression of MGMT has been associated with TMZ resistance. Clinical studies have observed a downward shift in MGMT methylation percentage from primary to recurrent stage tumors. However, the evolutionary processes driving this shift, and more generally the emergence and growth of TMZ-resistant tumor subpopulations, are still poorly understood. Here we develop a mathematical model, parameterized using clinical and experimental data, to investigate the role of MGMT methylation in TMZ resistance during the standard treatment regimen for GBM (surgery, chemotherapy and radiation). We first find that the observed downward shift in MGMT promoter methylation status between detection and recurrence cannot be explained solely by evolutionary selection. Next, our model suggests that TMZ has an inhibitory effect on maintenance methylation of MGMT after cell division. Finally, incorporating this inhibitory effect, we study the optimal number of TMZ doses per adjuvant cycle for GBM patients with high and low levels of MGMT methylation at diagnosis.

## 1 Introduction

Glioblastoma multiforme (GBM) is an aggressive form of brain cancer with poor prognosis. Typically, GBM patients are treated with surgical resection followed by radiation therapy and chemotherapy with the oral alkylating agent temozolomide (TMZ). This standard regimen results in median survival of only 15 months and a two-year survival rate of 30% [36]. The effectiveness of TMZ is impacted by the methylation status of the promoter for DNA repair protein O-6-Methylguanine-DNA Methyltransferase (MGMT). Clinical studies have linked epigenetic silencing of the MGMT gene via promoter methylation with greater sensitivity to TMZ and improved patient prognosis [20, 19], whereas resistance to TMZ has been associated with increased expression levels of MGMT [20, 42, 24].

Studies have compared MGMT promoter methylation in newly diagnosed tumors to matched recurrence samples following TMZ treatment [6, 37, 22, 11]. These studies provide evidence of a downward shift in the MGMT promoter methylation percentage during treatment. For example, in [6], 8 of 13 patients transitioned from an MGMT-methylated primary tumor to an unmethylated recurrent tumor after treatment, and in [37], it was reported that 10 of the 13 patients switched from a methylated primary tumor to an unmethylated recurrent tumor. In [11], authors observed that 39.1% of pretreatment GBM and 5.3% of recurrences were promoter methylated, in addition to an observed increase of MGMT activity in the recurrences. In [22] 15 of 18 recurrence samples displayed higher MGMT expression than matched primary samples. However, it is unclear whether this transition from methylated to unmethylated recurrent tumors is due to TMZ actively influencing the methylation status of MGMT, as some have hypothesized [9, 24, 6], simply a result of evolutionary selection for a more drug-tolerant phenotype, or some combination of both processes. We strive to understand this question by modeling the evolutionary processes driving this shift.

Previous works have mathematically modeled of the response of glioblastoma to treatment. In [26] the authors model chemotherapeutic delivery to brain tumors using a two compartment catenary model. In [35], a spatio-temporal model allowing for GBM patient-specific TMZ optimization is developed. The model in [5] explores the interactions between rapidly proliferating GBM cells and a dormant cell population. The effect of fractionated radiation dosing on GBM is studied using the linear-quadratic model [14, 8, 41, 3]. Powathil and colleagues consider a spatio-temporal brain tumor model including effects from both radiotherapy and chemotherapy in [32]. Patient-specific models of glioblastoma are developed in [33, 13] to predict patient response to radiotherapy and to determine optimal dosing strategies. Many more mathematical modeling efforts focusing on glioblastoma growth and therapy response are reviewed in [18]. Mathematical models have also been developed to describe the process of DNA methylation changes in cells [40, 30, 16, 34, 31]. For example, Otto and Walbot introduced the first model describing methylation in terms of maintenance and *de novo* methylation [30]; a similar model in a continuous-time framework was developed in [31]. We consider a discrete-time Markov chain version of Otto and Walbot’s methylation model, presented by Sontag and colleagues in [34].

Here, we develop and parameterize a stochastic model of the evolutionary dynamics driving GBM response to standard treatment. We incorporate a variant of the methylation model in [34] to investigate the role of MGMT promoter methylation in TMZ resistance. In particular, we focus on the specific roles of three major DNA methyltransferases (DNMT1, DNMT3a, and DNMT3b) within the methylation process, illustrated in Figure 1a. DNMT1 is responsible for ‘maintenance methylation,’ in which patterns of methylation in the original parental DNA are preserved in the replicated DNA. DNMT3a and DNMT3b are responsible for *de novo* methylation, in which unmethylated sites in the parental DNA become methylated in the replicated DNA [23, 10, 38]. By incorporating detailed mechanisms of maintenance and *de novo* methylation, driven by DNMT1, DNMT3a, and DNMT3b, within an evolutionary model of GBM treatment response, this work provides unique insight regarding the dynamics of MGMT methylation and GBM recurrence.

**Figure 1:**
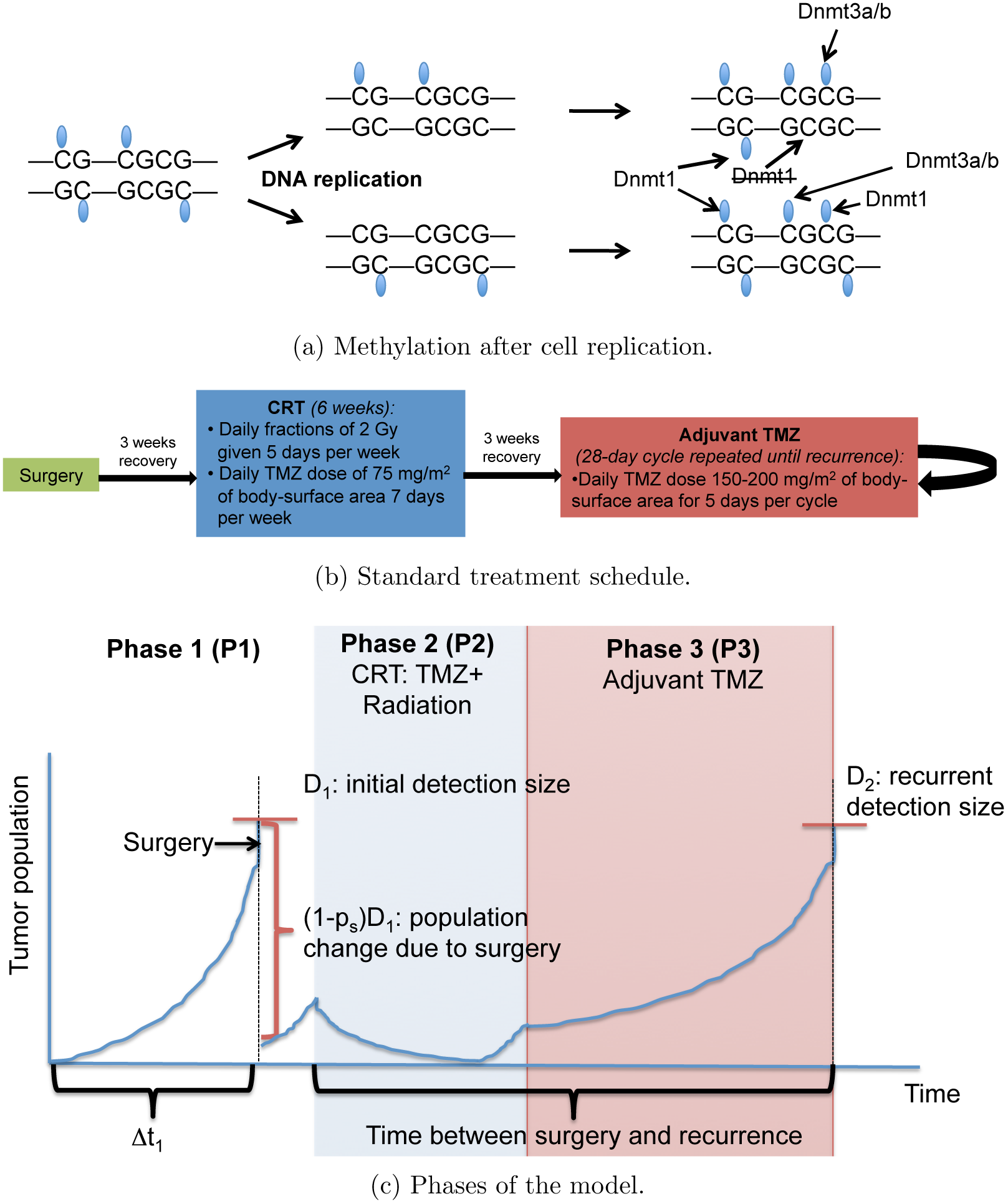
Diagram (a) illustrates a portion of a DNA molecule splitting during replication and the role of the DNA methyltransferases DNMT1 and DNMT3a/b. Notice that DNMT1 methylates the sites in the new strand that were methylated in the parental strand. As this process is not perfect, some sites can be missed. Dnmt3a/b methylates new sites that were not previously methylated in the top strands of the upper and lower molecules. This figure is similar to a figure in [34]. Figure (b) describes the standard treatment schedule for GBM [36]. Figure (c) depicts the three phases of the model; P1 consists of the tumor growth prior to detection and surgery, P2 denotes the concurrent radiation and chemotherapy (CRT) phase of treatment, and P3 refers to the adjuvant chemotherapy following CRT.

The outline of this paper is as follows: Section 2 describes the framework of the mathematical model, and Section 3 details the clinical and experimental data we collected and used to parameterize the model. In Section 4 we present our findings regarding methylation changes in tumors during therapy and optimal TMZ dosing strategies. We summarize these results and discuss future directions in Section 5.

## 2 Mathematical model

We develop a stochastic model describing the evolutionary dynamics of GBM response to standard treatment. We utilize a multi-type, continuous-time birth-death process model (see, e.g. [2]) in which each cell waits an exponential amount of time before division or death, governed by its birth and death rates. The model consists of three GBM cellular subtypes: Type-1, with fully methylated MGMT promoters; Type-2, with hemimethylated MGMT promoters; and Type-3, with unmethylated MGMT promoters. The type-1 cells are TMZ-sensitive, and type-2 and type-3 cells are considered TMZ-resistant since they both possess the ability to repair the lesion created by TMZ. Let *X*_1_(*t*)*, X*_2_(*t*), and *X*_3_(*t*) denote the number of type-1, type-2, and type-3 cells, respectively, at time *t*. Cellular birth and death rates vary during treatment with TMZ and radiation and are estimated with experimental data (see section 1 of the Supplement). The TMZ-resistant cells, *X*_2_(*t*) and *X*_3_(*t*), are assumed to have the same birth and death rates.

Conversions also occur between cell-types are driven by methylation/demethylation on the MGMT promoter immediately after cell division. To model these processes, we utilize a variant of the description in [34] to describe maintenance and *de novo* methylation at a CpG site. This underlying methylation model feeds into our population-level branching process model via the rates of conversion between the cellular subtypes. More specifically, let *ρ* be the probability of maintaining methylation for any given CpG site after replication, i.e the probability that DNMT1 methylates a CpG dyad after replication, conditioned on the event that the site was methylated before replication. Let *ν* be the probability of *de novo* methylation, i.e. the probability that DNMT3a or DNMT3b methylates any CpG site that is unmethylated immediately following DNA replication. Inspired by [34], we derive the following offspring distributions for each of the three cell types, conditioned on cell division. In these distributions, *p_i_*(*x, y, z*) refers to the probability that a type-*i* cell will produce *x* type-1 cells, *y* type-2 cells, and *z* type-3 cells after replication. For ease of notation, let

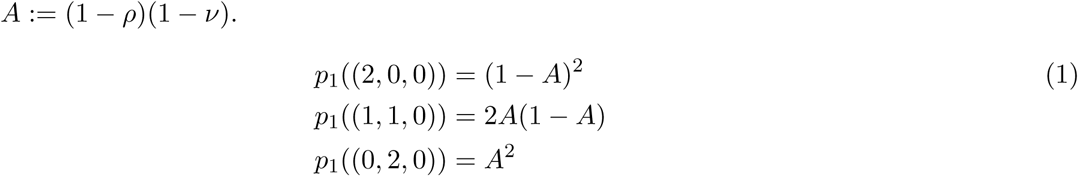

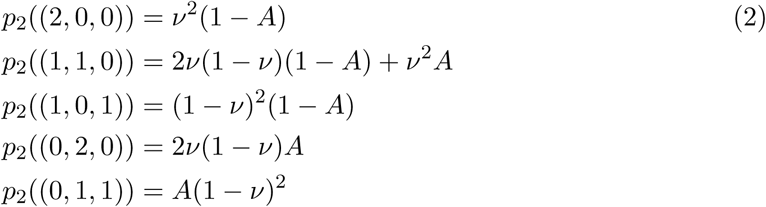

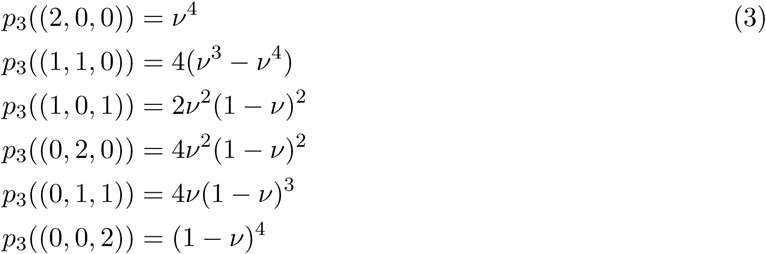

See Supplementary Information Figure 1 depicting the set of possible birth events. Note that a methylated dyad produces two hemimethylated dyads when the DNA strands split during replication, and those sites remain hemimethylated if the site without methylation is not methylated by DNMT1 or DNMT3a/b immediately following replication. Hence, the probability that each dyad remains hemimethylated is *A* = (1 *−ρ*)(1 *−ν*), and consequently the probability of producing two hemimethylated dyads, i.e. two type-2 cells, is *A*^2^. Conversely, the probability that one of those hemimethylated sites becomes fully methylated is 1 *− A*, so the probability of producing two fully methylated (type-1) cells is (1 *− A*)^2^, and the probability of producing one type-2 and one type-1 cell is *A*(1 *− A*). The offspring distributions for type-2 and type-3 cell replication can be verified similarly upon inspection, using the idea that an unmethylated dyad produces two unmethylated dyads during replication, and each CpG site within these dyads can only be methylated with DNMT3a/b, i.e via *de novo* methylation. Later, as we investigate the potential impact of TMZ on the methylation processes, we allow the *de novo* and maintenance probabilities to vary in the presence of the TMZ.

The binary stratification of tumors into ‘MGMT-methylated’ or ‘MGMT-unmethylated’ in clinical literature requires some clarification, since methylation status can vary between tumor cells and between CpG sites in the same genic promoter region. Typically the percentage of methylated cells is determined for a small subset of the CpG sites in the MGMT promoter region and averaged across the sites. Then a threshold, which may vary widely between studies, is used to stratify tumors into ‘methylated’ vs ‘unmethylated’ status. Due to substantial variation in the definition of this threshold between studies, here we model quantitative changes in methylation percentage on a representative CpG site, rather than imposing a binary stratification.

The model describes three phases of tumor development and standard GBM treatment. Phase I (P1) consists of tumor growth before detection, surgery, and a three-week recovery. Phase 2 (P2) consists of concurrent radiotherapy and chemotherapy (CRT) for six weeks, followed by a three-week recovery. During CRT, daily radiation fractions of 2 Gy are given five days per week, and 75 mg of TMZ per square-meter of body-surface area are administered daily. Phase 3 (P3) consists of repeated 28-day cycles of adjuvant chemotherapy (five daily doses of 150-200 mg/m^2^ TMZ, followed by a 23-day recovery) until tumor recurrence. Further details are described in [36]. Schematics of the standard GBM treatment schedule and the three phases are provided in Figure 1. Below we describe the adaptation of the branching process dynamics during each phase.

### Phase I: Pretreatment, surgery, recovery

In the absence of treatment, the intrinsic birth rates of untreated methylated (type-1) and hemi/unmethylated (type-2/ type-3) cells are *b*_1_ and *b*_2_ per day, respectively, and their death rates are *c*_1_ and *c*_2_. The parameters of the model are determined using experimental and clinical data (see section 1 of the supplement); a summary of the baseline parameter set is provided in Table 1 of the Supplement. Once the tumor population reaches a detection size threshold *D*_1_, we model surgical resection of the tumor by removing *p_s_* percent of the total cells, chosen proportionally for each subtype. During the three-week recovery period, the initial birth and death rates drive the regrowth of the tumor.

### Phase II: TMZ, radiation for six weeks and recovery

In P2, the tumor undergoes concurrent radiotherapy and chemotherapy for 6 weeks. The standard schedule for radiotherapy is a daily fraction of 2 Gy, given five days per week, on Monday through Friday. In addition, the tumor is treated every day with 75 mg of TMZ per square-meter of body-surface area. Since TMZ is a cytotoxic treatment, we model its impact by increasing the death rates of the tumor cells, denoted *c*_1_ and *c*_2_. Let *g*_1_(*t*)*, g*_2_(*t*) be the additional death rate due to TMZ treatment for type-1 and type-2/type-3 cells, respectively. These rates depend on the current TMZ concentration level and are determined from experimental and pharmacokinetic data, detailed in Supplementary Information.

The cytotoxic effect of radiotherapy is modeled using the standard linear-quadratic (L-Q) model [41]. Here, radiosensitivity parameters *α, β* are used to account for toxic lesions to DNA and misrepair of repairable damage to DNA, respectively [15]. Under the L-Q model, the probability of cell survival at time *t* is dependent on the radiation dose at time *t*, *D*(*t*): *S*(*t*) = *e*^*−αD*(*t*)*−βD*(*t*)^2^^. At the time *t* of each radiation dose, we remove (1*−S*(*t*))*X*_1_ (*t*) type-1 cells and (1 *− S*(*t*))*X*_2_(*t*) type-2 cells. During the three-week recovery, cellular birth and death rates revert to those used in the pretreatment phase. Given data constraints, we ignore differences in radiosensitivity between type-1 and type-2/3 cells.

### Phase III: Adjuvant TMZ

During P3, adjuvant chemotherapy is administered to the tumor. In this phase, the additional death rates *g*_1_(*t*) and *g*_2_(*t*) due to chemotherapy reflect five daily TMZ doses of 150-200 mg/m^2^, followed by 23 days off. This 28-day cycle repeats until tumor recurrence, which occurs when the tumor population size reaches the threshold *D*_2_.

## 3 Experimental and clinical data

### Experimental setup

We performed experiments on PDX cell lines to investigate the differential impact of TMZ on the growth kinetics of MGMT-methylated and unmethylated GBM cells. In these *in vitro* experiments, plates of GBM6 cells were treated at eight concentrations of TMZ (including DMSO) in triplicate, and live and dead cell counts were collected via MTT and trypan blue assays after 8 days of exposure. The average number of live cells for each TMZ concentration is displayed in Figure 2b. Figure 2c plots the average proportion of live cells from the sum of live and dead cells after 8 days of exposure, for various TMZ doses. The frequency of cells expressing MGMT, assessed in each group after eight days, is displayed in Figure 2a. This data is used in section 1 of the supplement to determine the cell viability for type-1 and type-2/3 cells, as a function of TMZ concentration.

**Figure 2:**
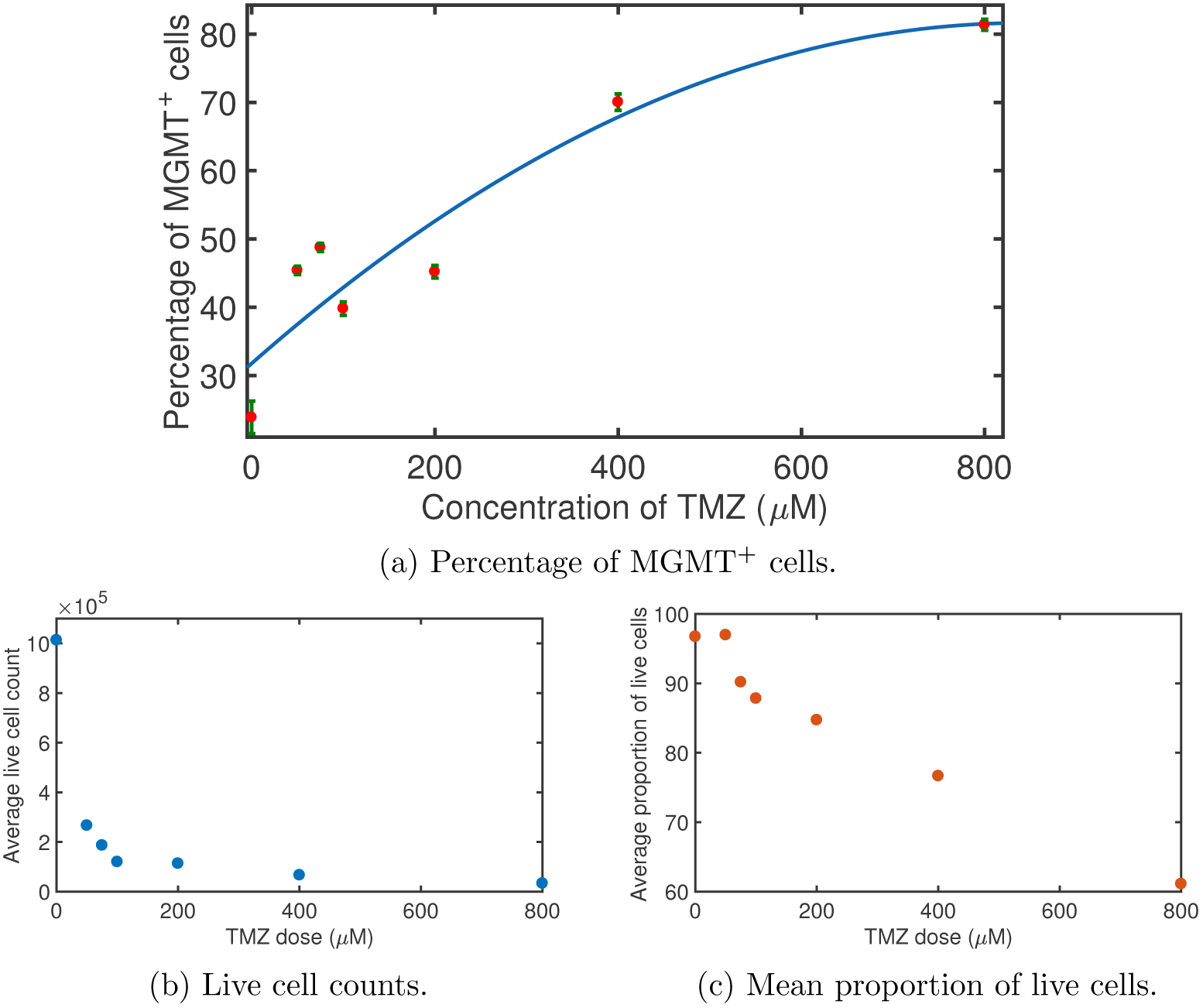
Plot (a) shows the percentage of cells expressing MGMT after 8 days of exposure to various concentrations of temozolomide (in *νM*), assessed using PDX experiments. The red dots in the plot denote average percentages of MGMT^+^ cells, and the error bars indicate the standard deviation. Plot (b) displays the average live cell counts collected after 8 days of exposure to TMZ, as a function of the concentration of TMZ exposure (in *μM*). Plot (c) shows the mean proportion of live cells, out of the total cells (live and dead cells) after 8 days of exposure, as a function of the concentration of the TMZ dose (in *μM*).

### Clinical data

Clinical data was also collected from a group of 20 adult patients that received the standard protocol described in [36]. Tumor radius size was collected from each patient at initial detection and at recurrence. This data is summarized in Figure 3a. Tumor growth in the absence of treatment was also tracked, resulting in net growth rate estimates for each patient, summarized in Figure 3c. Using the patient data and a reaction-diffusion model described in [12], we obtained an average net growth rate estimate of *λ*= 0.0897/cell/day before treatment. Patient data describing the tumor radius size after surgery is displayed in Figure 3c. The clinical data is used in section 1 of the Supplement to characterize the tumor size at detection and after surgery within the model, and the intrinsic cellular growth rates in the absence of treatment.

**Figure 3:**
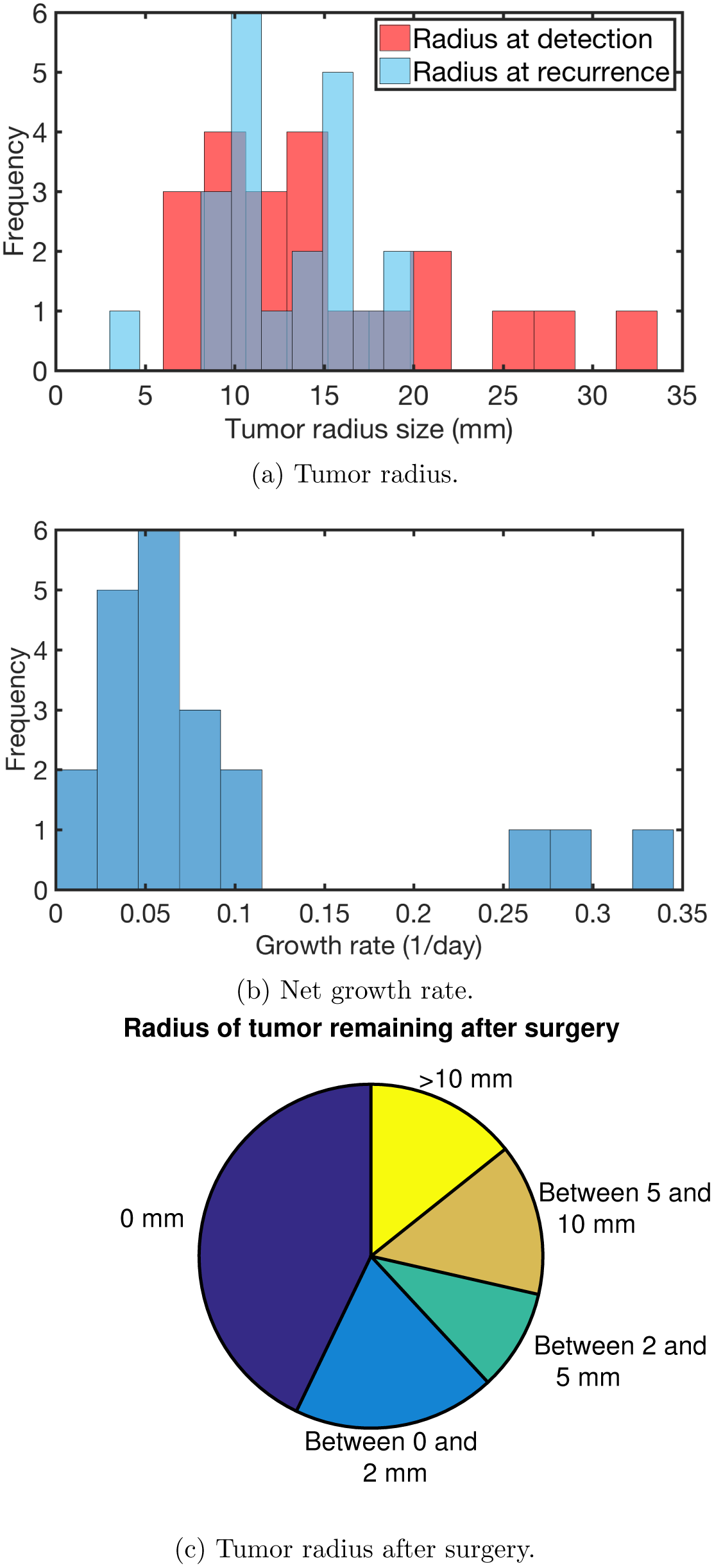
Clinical data from GBM patients undergoing standard regimen (*n* = 21). Histograms of (a) tumor radius at detection and recurrent tumor radius (mm), (b) the overall net growth rate (1/day) before treatment, and (c) a pie chart depicting the radius of tumor remaining after surgery (mm).

## 4 Results

To demonstrate the model dynamics, we first provide a single sample path simulation of the model in Figure 4a, showing the type-1 (methylated) and total population sizes during the standard treatment regimen. Figure 4b shows the distribution of recurrence times from a computational experiment with 100 Monte Carlo simulations; the median recurrence time is 213.8 days, roughly consistent with clinical data in [29], reporting a median recurrence time of 191 days.

**Figure 4:**
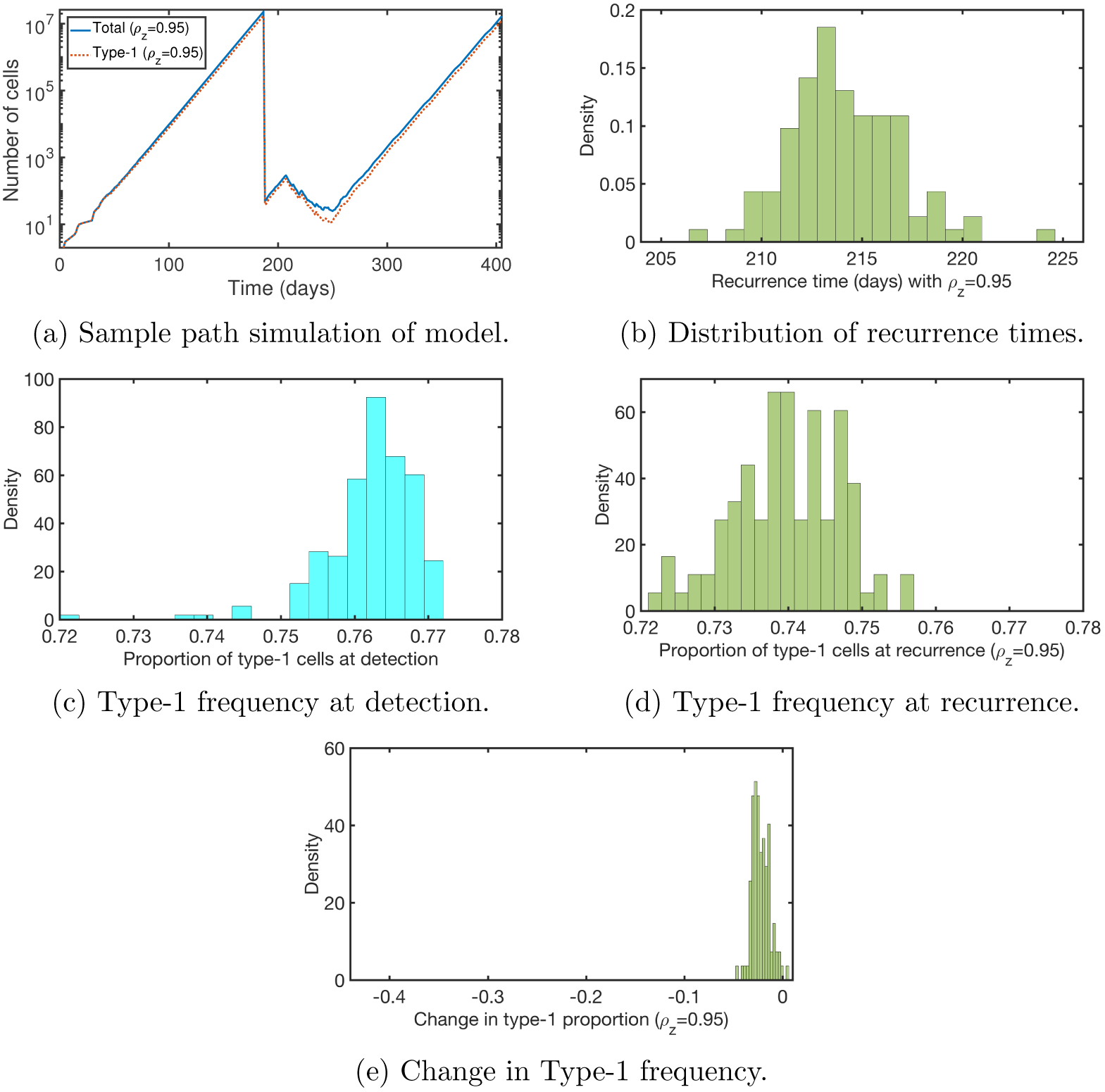
Simulation results with no TMZ impact on methylation rates. Plots of (a) one sample path simulation of the model, (b) the distribution of recurrence times in a computational experiment with 100 samples, (c) the distribution of methylation percentage at the time of detection, (d) the distribution of methylation percentage at the time of recurrence, and (e) the distribution of change in methylation percentage between detection and recurrence. All parameters are set as described in Table 1 of the Supplement.

### Selection alone does not explain methylation shift in recurrent tumors

Using the parameter settings obtained from the calibration described in supplementary information, we next examined the relative methylation percentages at diagnosis and recurrence. Figures 4c and 4d show the distribution of methylation percentages found at these times. We observe that the average proportions of methylated (type-1) cells to total cells at recurrence and diagnosis are roughly the same. The distribution of the change in methylation percentage between detection and recurrence is depicted in Figure 4e. The slight reduction in overall methylation percentage suggests that evolutionary selection alone cannot account for the significant reduction in methylation observed in the clinical studies described in section 1.

### TMZ inhibition of maintenance methylation results in downward methylation shift

We next investigated the hypothesis that TMZ actively impacts the cellular methylation processes, driving the methylation downshift in recurrent tumors. In particular, we investigated the possibility that TMZ may decrease the amount of time spent in the type-1 (methylated) state and increase time spent in type-2/3 states. This may result from a decrease in either the *de novo* methylation probability *ν* or the maintenance methylation probability *ρ*. Note that for this investigation, the parameters *ν, ρ* will deviate from their baseline values only during TMZ treatment periods; we denote the parameters during TMZ treatment as *ν_z_, ρ_z_*.

To investigate the effects of changing the *de novo* methylation probability, *ν_z_*, in the presence of TMZ, we first note that the lowest possible value of *ν_z_* is 0, representing no *de novo* methylation events. When we let *ν_z_* = 0, we observe a modest decrease in the expected methylation percentage between detection and recurrence, changing by about 7%. Figures 5a and 5b show the distribution of methylation percentages at the time of recurrence and the distribution of change in methylation percentage between detection and recurrence, respectively, when *ν_z_* = 0. Thus, a significant drop in methylation percentage after TMZ treatment cannot be attributed to an inhibitory impact on *de novo* methylation.

**Figure 5:**
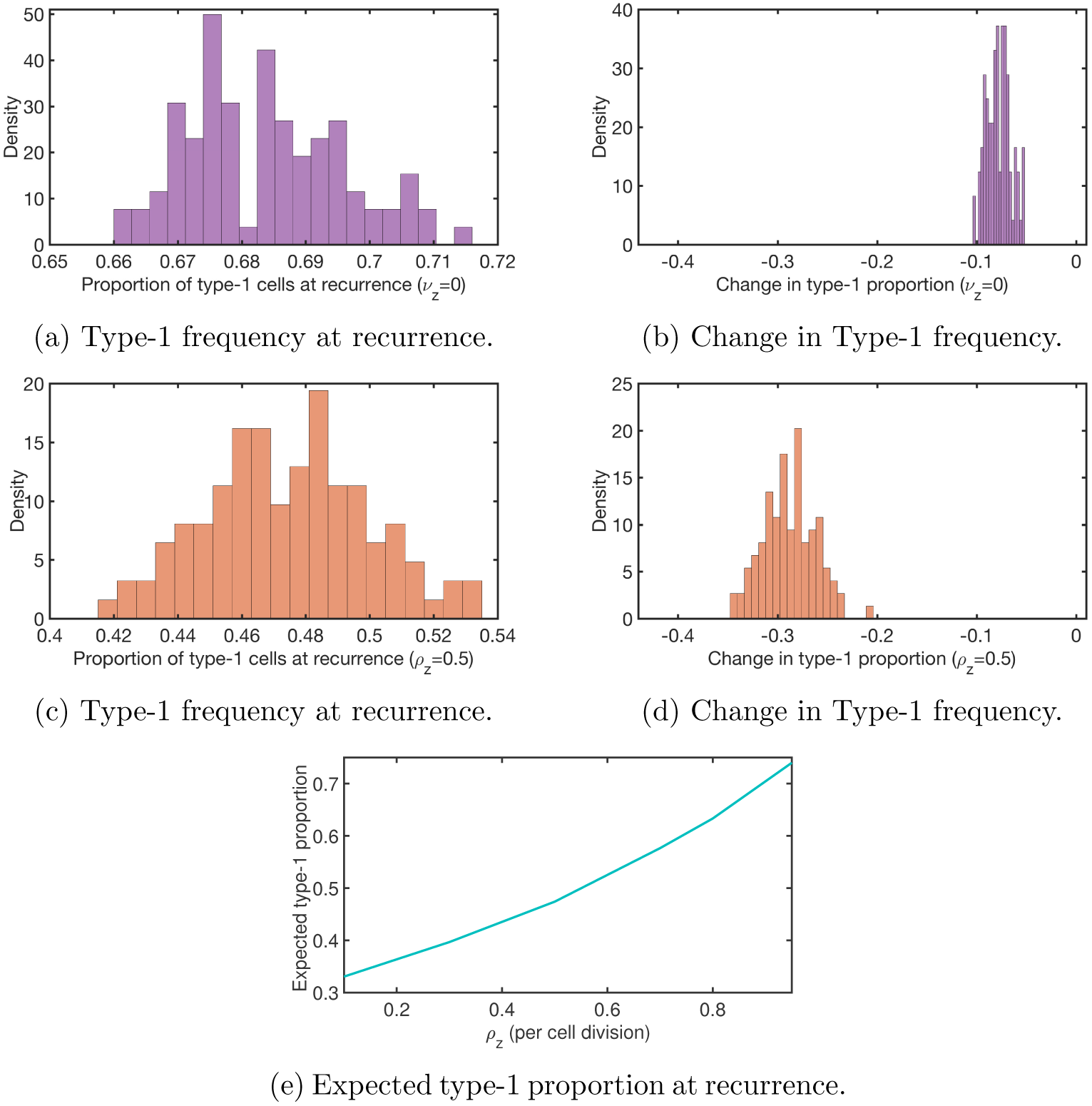
Simulation results with TMZ inhibition of methylation rates. Plots of (a) the distribution of methylation percentage at the time of recurrence when *ν_z_* = 0, (b) the distribution of change in methylation percentage between detection and recurrence when *ν_z_* = 0, (c) the distribution of methylation percentage at the time of recurrence when *ρ_z_* = 0.5, (d) the distribution of change in methylation percentage between detection and recurrence when *ρ_z_* = 0.5, and (e) expected proportion of type-1 cells at recurrence under the standard treatment schedule, as a function of the maintenance methylation probability, *ρ_z_*. Non-varying parameters are set to the baseline values described in Table 1 of the Supplement.

We next investigated the impact of decreasing the probability of maintenance methylation, *ρ_z_*, during chemotherapy. Figure 5c displays the type-1 frequency at the time of recurrence when *ρ_z_* = 0.5, reduced from the baseline value of *ρ* = 0.95, and figure 5d shows the change in methylation percentage between detection and recurrence. We observe that there is a much more significant decrease in methylation in this case than in the case when TMZ does not impact methylation rates (compare Figures 5d and 4e). Figure 5e displays the expected proportion of type-1 cells at recurrence, as a function of *ρ_z_*. For smaller values of *ρ_z_*, the proportion of type-1 cells after treatment decreases substantially from a mean methylation percentage of 0.762 at detection. Thus, TMZ inhibition of maintenance methylation, but not *de novo* methylation, can explain the clinically observed downward shift in methylation. In section 2 of the supplement, we show that this claim is robust to model parameter variability.

### Optimization of adjuvant TMZ schedule to minimize expected tumor size

We next used the model to investigate the optimal number of TMZ doses during Phase III, the adjuvant chemotherapy phase, to minimize the expected tumor size after 4 cycles of treatment. In the standard treatment schedule, five TMZ doses of 150-200 mg/m^2^ are administered daily at the beginning of each 28-day cycle. Let *n* denote the number of TMZ doses in a single 28-day cycle. We vary *n* in order to determine the number of doses and dose level that minimizes the number of total cells remaining after 4 adjuvant cycles. Let *Z*(*n*) denote the TMZ concentration level per dose, in mg/m^2^, when *n* doses are administered per cycle. Each dose concentration is set at *Z*(*n*) = 1000*/n* for varying values of *n*, where 0 *< n ≤* 28, so that the cumulative TMZ dosage per cycle does not exceed 1000 mg/m^2^. Based on our previous investigations, the maintenance methylation probability in the presence of TMZ, *ρ_z_*, is assumed to be 0.5.

Mean calculations for each cell-type, provided in section 3 of the supplement, are used to determine the *n* that minimizes the expected tumor size after 4 cycles. Figures 6a and 6b show the mean tumor size, number of fully methylated cells (type-1), and cells that are not fully methylated (type-2 and type-3) when *ρ_z_* = 0.5. In this case, the optimal number of doses per cycle, i.e. the number that results in the smallest mean tumor population after 4 cycles, is *n* = 6, with *Z*(6) = 166.67 mg/m^2^. This is close to the standard administered dose during adjuvant chemotherapy, and we see a small difference in the expected tumor size when 5 vs 6 doses are administered. Hence, our model suggests that the standard dosing schedule is a reasonable, though not optimal, protocol for highly methylated tumors at diagnosis.

**Figure 6:**
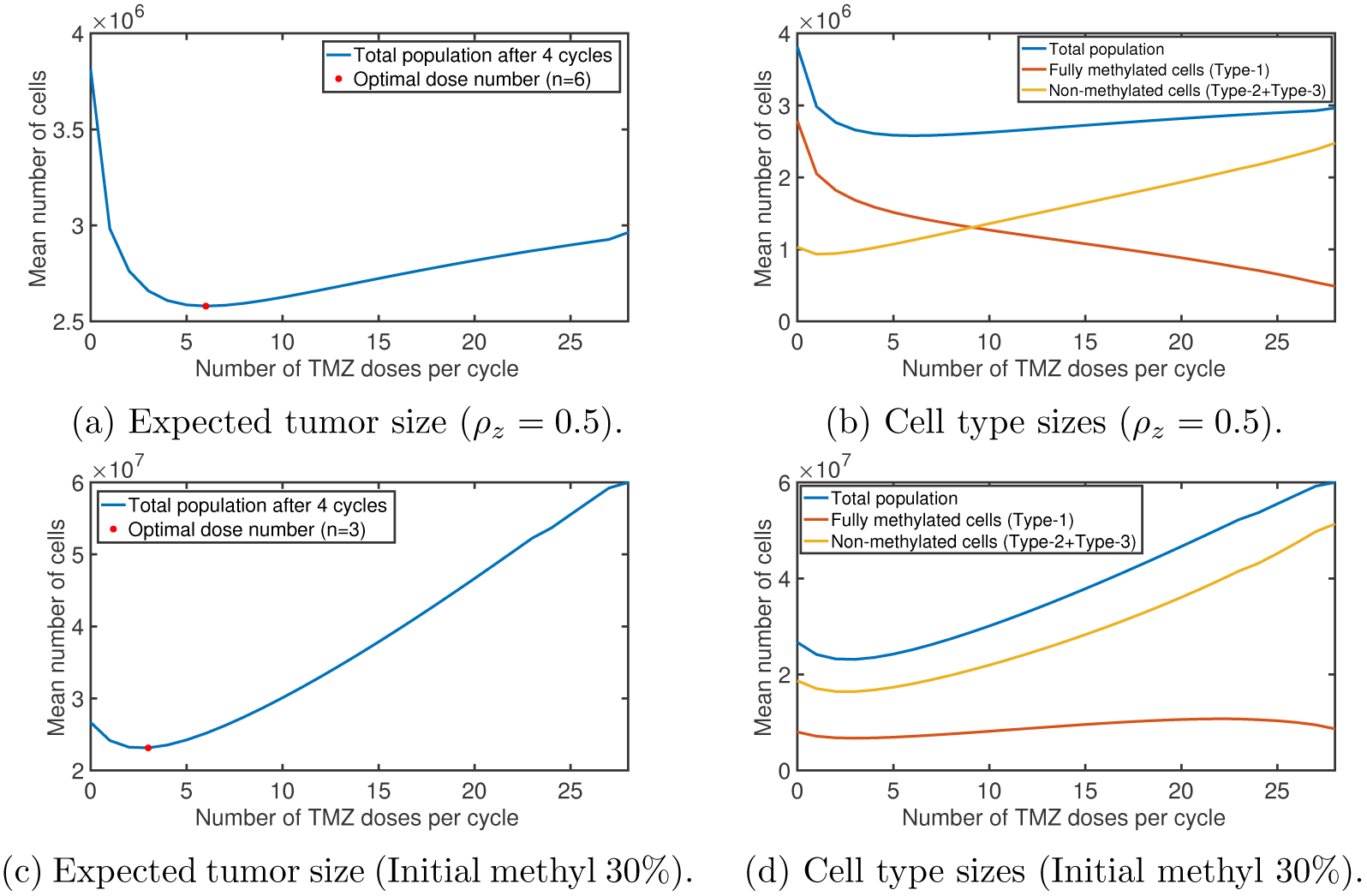
Adjuvant TMZ optimization comparison between tumors with high and low levels of methylation at diagnosis. Plots of (a) the mean tumor population size and (b) the mean total, type-1, and type-2/3 cell population size when *ρ_z_* = 0.5, and (c) the mean tumor population size and (b) the mean total, type-1, and type-2/3 cell population size, when the expected methylation proportion at diagnosis is 0.3. The mean cell populations are calculated after 4 adjuvant chemotherapy cycles, as a function of the number *n* of doses in one cycle during P3. We use the standard set of parameters. In (a) and (c), we also plot the optimal number of TMZ doses (*n* = 6 and *n* = 3, respectively) and the corresponding tumor size in red.

We also used the model to investigate the optimal adjuvant TMZ schedule for tumors with lower methylation percentages at diagnosis. To this end, we first identified the combination of birth rates (*b*_1_ = .0569 day^*−*1^ and *b*_2_ = 0.1276 day^*−*1^) that satisfied the net growth rate constraint and led to 30% methylation at detection. Figures 6c and 6d display plots of the mean number of total, fully methylated (type-1), and non-methylated (type-2 and type-3) cells, as functions of the number of doses per cycle. We observe that the tumor is dominated by non-methylated cells for all *n*, and the large population of TMZ-resistant cells makes a large number of TMZ doses less effective. Additionally while *n* = 3 is the optimal dose number in this case, it is not significantly more beneficial than no adjuvant TMZ treatment. Such behavior is consistent with clinical observations; the study in [20] found that unmethylated tumors treated with radiotherapy and the standard TMZ regimen had a median overall survival of 12.7 months, versus a median overall survival of 11.8 months for those receiving only radiotherapy. Thus, our model suggests that tumors with low levels of methylation at diagnosis may be better served by alternative therapies, such as O^6^-benzylguanine discussed in [1], that can be used in combination with TMZ, to counter TMZ’s impact on the methylation process.

## 5 Discussion

In this work we developed a mathematical model integrating a mechanistic description of MGMT promoter methylation/demethylation with the evolutionary dynamics of GBM tumor progression during standard treatment. We investigated several possible causes for the clinical observed drop in methylation percentage between primary and recurrent tumor stages. Our results indicate that this clinically observed drop in methylation between diagnosis and recurrence cannot be explained by evolutionary selection, suggesting that TMZ may have an active role in altering methylation processes. Investigating this further, we found that TMZ inhibition of the maintenance methylation results in a sizable reduction in expected methylation percentage at recurrence, consistent with clinical results.

The precise mechanism by which TMZ may contribute to MGMT demethylation is unclear, but experimental studies suggest this may involve the activation of the protein kinase C (PKC) signaling pathway. In [4] it was demonstrated that alkylating drugs similar to TMZ led to an increase in MGMT expression and in PKC activity. In [25], the authors discovered that a number of PKC isoforms induce the attachment of a phosphoryl group to DNMT1. Further testing on the specific isoform PKC*ζ* showed that cells with a high expression of both PKC*ζ* and DNMT1 exhibited a significant reduction in methylation; this was not the case in cells with a high expression of PKC*ζ* or DNMT1 alone. This suggested that the methylation reduction results from the phosphorylation of DNMT1, driven by PKC*ζ*; another study in [21] confirms that the phosphorylation of DNMT1 is associated with hypomethylation of gene promoters. Hence, experimental studies suggest that TMZ may contribute to MGMT demethylation by activating the PKC signaling pathway in GBM cells, leading to the phosphorylation of DNMT1, thereby inhibiting maintenance methylation within the affected cells, as our model suggests.

Incorporating the proposed TMZ effect, we used the model to find the optimal number of TMZ doses administered during adjuvant chemotherapy. The number of daily TMZ doses administered during each cycle was varied while maintaining the same cumulative dosage per cycle, to determine the dose number that minimizes the mean tumor population after 4 adjuvant cycles. We determined an optimal TMZ dosing schedule of 6 daily doses of 166.67 mg/m^2^, followed by 22 days off. The standard schedule of 5 daily doses/cycle is nearly optimal, resulting in a slightly larger mean tumor size after 4 cycles. We also investigated the optimal adjuvant chemotherapy schedule for a tumor with a low methylation percentage at diagnosis. Receiving three larger doses of TMZ is optimal in this case, but does not provide a significant benefit over the absence of any adjuvant TMZ treatment. This observation is consistent with clinical results comparing the benefit of both radiotherapy and chemotherapy versus radiotherapy alone for unmethylated primary tumors. Therefore, our model suggests that for primarily unmethylated tumors it may be more beneficial to administer, in combination with TMZ, a therapy that can stimulate MGMT methylation within the tumor.

A limitation of our model is that we do not incorporate the diffuse nature of GBM tumors. We also assume that all hemimethylated and unmethylated GBM cells behave with the same intrinsic growth rates and that MGMT methylation status does not affect radiosensitivity. Note that a few studies have suggested that there may be a phenomenon of MGMT depletion after extended exposure to TMZ, in an attempt to explain observed differences in dose-dense TMZ treatment and the standard TMZ regimen [7, 39]. If this phenomenon occurs, it would make TMZ more effective in tumors with low methylation levels. However, other studies have found no conclusive differences in dose-dense TMZ regimens and the standard TMZ regimen for all GBM [27, 17]. Thus, we did not incorporate a mechanism for MGMT depletion in our model. In future work, we plan to investigate the hypothesis that the oncogenic IDH1 mutation drives increased methylation in gliomas [28].

## 6 Acknowledgements

K.S. and J.F. were supported under National Science Foundation grant DMS-1349724. K.S and K.L. were supported under National Science Foundation grant CMMI-1362236. R.C.R. was supported by the National Cancer Institute of the National Institutes of Health under award number P30CA033572. A.U.A. was supported by the National Institute of Neurological Disorders and Stroke grant 1R01NS096376 and the American Cancer Society grant RSG-16-034-01-DDC. We thank Dr. C. David James for providing all the patient derived xenografts lines. The content is solely the responsibility of the authors and does not necessarily represent the official views of the funding organizations above.

